# Phospho-regulation of the Shugoshin - Condensin interaction at the centromere in budding yeast

**DOI:** 10.1101/2019.12.16.877894

**Authors:** Galal Yahya Metwaly, Yehui Wu, Karolina Peplowska, Jennifer Röhrl, Young-Min Soh, Frank Bürmann, Stephan Gruber, Zuzana Storchova

## Abstract

Correct bioriented attachment of sister chromatids to mitotic spindle is essential for chromosome segregation. The conserved protein shugoshin (Sgo1) contributes in budding yeast to biorientation by recruiting the protein phosphatase PP2A-Rts1 and the condensin complex to centromeres. Using peptide prints, we identified a Serine-Rich Motif (SRM) of Sgo1 that mediates the interaction with condensin and is essential for centromeric condensin recruitment and the establishment of biorientation. We show that the interaction is regulated via phosphorylation within the SRM and we determined the phospho-sites using mass spectrometry. Analysis of the phosphomimicking and phosphoresistant mutants revealed that SRM phosphorylation disrupts the shugoshin – condensin interaction. We present an evidence that Mps1, a central kinase in the spindle assembly checkpoint, directly phosphorylates Sgo1 within the SRM to regulate the interaction with condensin and thereby condensin localization to centromeres. Our findings identify novel mechanisms that control shugoshin activity at the centromere in budding yeast.

**Author summary:** Proper chromosome segregation in eukaryotes is ensured through correct attachment of the spindle microtubules to the centromeric chromosomal regions. The attachment is mediated via the multimolecular proteinaceous complex called kinetochore and precisely regulated. This enables the establishment of so called bioirentation, when each sister chromatid is attached to microtubules emanating from opposite spindle poles. Shugoshin (Sgo1) is a conserved centromeric protein that facilitates biorientation through its interactions with the protein phosphatase PP2A/Rts1, chromosome passanger complex and centromeric condensin. Here, we identified a serin-rich motif that is required for the interaction of shugoshin with the condensin complex. We show that loss of this region impairs condensin enrichment at the centromere, chromosome biorientation, segregation as well as the function of the chromosome passanger complex in the error correction. Moreover, the interaction is phosphoregulated, as phosphorylation of the serin-rich motif on Sgo1 disrupts its interaction with condensin. Finally, we show that the conserved spindle assembly checkpoint kinase Mps1 is responsible for this phosphorylation. Our findings uncover novel regulatory mechanisms that facilitate proper chromosome segregation.

## Introduction

Biorientation of sister chromatids relies on two major processes. First, spindle and kinetochore geometry facilitates the capture of sister kinetochores (KT) by microtubules (MTs) emanating from the opposite spindle poles (1, 2). Second, destabilization of incorrect, tension-less interactions between KTs and MTs allows error correction (3), while the spindle assembly checkpoint (SAC) holds the progress through mitosis until bioriented sister kinetochore attachments are achieved (4). Several proteins, whose activity must be tightly regulated and coordinated, are required for these processes. Among these proteins are so-called shugoshins, a poorly conserved family of proteins with an important function in establishment of biorientation in mitosis and meiosis (5). A body of evidence determined shugoshins as protectors of centromeric cohesion from separase cleavage during meiosis (e.g. (6)). Shugoshins also prevent cohesion loss at the centromere during mammalian mitosis, although via a distinct mechanism that affects the prophase pathway (7). These activities are carried out through the recruitment of a protein phosphatase 2A (PP2A-Rts1 in budding yeast, PP2A-B56 in mammalian cells) that reduces cohesin removal via its dephosphorylation (8). Shugoshin proteins also recruit additional proteins to the centromeres, such as the chromosome passenger complex (CPC), MCAK (mitotic centromere-associated kinesin)/Kif2A kinesin motor, and condensin (5). The activity of shugoshins thereby facilitates centromeric conformation, establishment of biorientation, tension sensing and correction of aberrant MT-KT attachments.

Budding yeast *Saccharomyces cerevisiae* encodes one Sgo1 variant. Similarly as in other species, Sgo1 localizes to the centromeric chromatin by binding to histone H2A phosphorylated on serine 121 (T120 in human) by the SAC kinase Bub1 (9). This interaction is mediated by a basic region, one of the two conserved regions of shugoshin proteins (Fig. 1A). Mps1, a central kinase in the SAC, is also required for the localization of shugoshin to the pericentromere in budding yeast (10). The interaction between Rts1, the regulatory subunit of PP2A, and Sgo1 is essential for the majority of known Sgo1 functions. Mutational analysis determined a region within the N-terminal coiled-coil domain of Sgo1 that is weakly conserved among species and mediates the interaction with Rts1(8) (11, 12) (13, 14). Sgo1 also contains an unusual D-box-related sequence motif near its C-terminus that directs its APC/cyclosome dependent degradation at the end of anaphase (Fig. 1A) (13). Another function of Sgo1 is to maintain centromeric enrichment of Ipl1/Aurora B, the kinase subunit of the CPC (9, 12, 14, 15), although the specific region of Sgo1 that is required for this function remains enigmatic. Additionally, Sgo1 recruits condensin to the budding yeast centromere (14, 16). There is only limited understanding of the nature of the interaction between shugoshin and condensin and its regulation. The Sgo1-condensin interaction is not dependent on DNA, suggesting that complex formation between Sgo1 and condensin does not require association with chromatin (16). Therefore, we hypothesized that there is a direct interaction between Sgo1 and subunits of the multi-subunit complex condensin. Moreover, given the importance of the condensin localization to the kinetochore for correct chromosome segregation, we hypothesized that this interaction might be regulated by spindle assembly checkpoint kinases.

**Figure 1.**
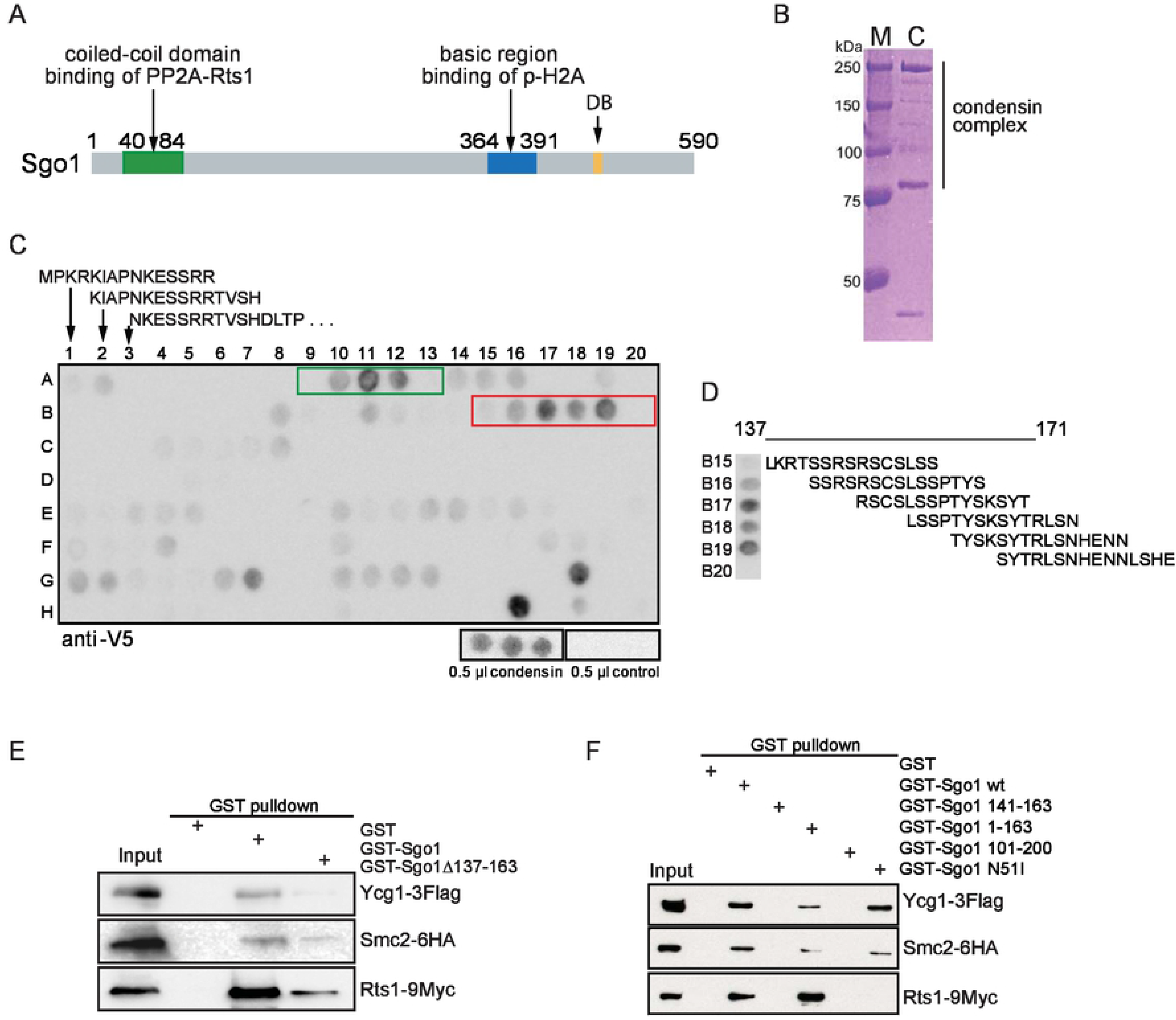
Identification of the motif of Sgo1 required for binding of the condensin complex. A) Schematic depiction of the known domains of Sgo1 in budding yeasts. DB – destruction box. B) The condensin complex was purified from budding yeast via pull down of a TwinStrep-tagged Smc2. M: Marker, C: purified condensin. C) Mapping of the binding sites of Sgo1 to the condensin complex using a peptide array. A1-H5: Sgo1 peptides, starting from position one, 15 aa, four aa overlap. H8 – H20: tags for antibody controls. Binding of the condensin complex with peptides A10, A11, A12 (amino acids from 37 to 59) and B16, B17, B18 (amino acids from 141 to 163) was considered as positive and validated in a secondary screen (Figure S1A, B). D) Binding of condensin to the Sgo1 peptides from positions B15 to B20. E) Yeast extracts were incubated with glutathione-coupled beads pre-treated with GST (line 2), GST-Sgo1wt (line 3) and GST-Sgo1 Δ137-163 (line 4), as well as F) GST-Sgo1 137-163 (line 4), GST-Sgo1 1-163 (line 5), GST-Sgo1 101-200 (line 6) and GST-Sgo1 N51I (line 7). The eluates were analyzed by immunoblotting with anti-FLAG (Ycg1), anti-HA (Smc2) and anti-MYC (Rts1) antibodies, respectively. Line 1: Input - yeast whole cell extract. For Coomassie stained gel of purified GST-tagged proteins see Fig. S1C.

To determine how Sgo1 recruits condensin to the centromeric region, we identified a region within Sgo1 that is essential for interaction with condensin *in vitro*. We demonstrate that a loss of the interaction motif impairs localization of condensin to the centromeres *in vivo*. The Sgo1 mutations that fail to recruit condensin to the centromeres also negatively affect sister chromatid biorientation and segregation. Finally, we determine that the interaction of Sgo1 with condensin is phosphoregulated. Based on our results we postulate that, in budding yeast, tightly regulated presence of condensin on centromeres is essential for correct chromosome segregation and is mediated via its interaction with Sgo1.

## Results

### Identification of the condensin-interacting region within Sgo1

To identify the regions responsible for the Sgo1 – condensin interactions, we prepared a peptide array of 15 amino acid peptides with four amino acids phase shifts covering the full length of Sgo1. The membrane with the spotted peptide array was then incubated with the condensin complex that was purified via the Smc2-TEV-HaloTag-TwinStrep tag from budding yeast (Fig. 1B). Interacting condensin complex bound to Sgo1 peptides was detected by far western via immunoblotting with an anti-V5 antibody that recognizes Smc4-V5-His6. We identified two putative regions of interaction between Sgo1 and condensin complex (Fig. 1C). To validate the results, we repeated the peptide print specifically for the identified regions at a higher resolution. As a control, we performed identical far western using the same purification procedure, but from an untagged yeast strain (Fig. S1A). This experiment confirmed the interaction of purified condensin with the Sgo1 peptides that covered an area from 137 to 163 amino acid (L_137_KRTSSRSRSCSLSSPTYSKSYTRLSN_163_) (Fig. 1D). To reflect its amino acid composition, we named this area Serine-Rich Motif (SRM). Other regions were not confirmed to interact with condensin in the secondary analysis (Fig. S1B).

To further corroborate the interaction, GST-fused Sgo1 and its mutants purified from *E. coli* (Fig. S1C) were used in pull down experiments to determine the interaction with condensin as well as with Rts1, the regulatory subunit of the phosphatase PP2A that binds the coiled-coil domain of Sgo1. We found that the full length Sgo1-GST pulls down Ycg1-3FLAG and Smc2-6HA, two subunits of the condensin complex, and Rts1-9Myc, as previously observed (Fig. 1E). Deletion of the SRM (aa 137-163) significantly reduced the interaction with condensin subunits (Fig. 1E). The Serine-Rich Motif alone as well as a larger fragment of Sgo1 containing SRM (101-200 aa) was not sufficient to pull down condensin subunits, while the N-terminal region of Sgo1, amino acids 1 – 163, interacted with both condensin and Rts1 (Fig. 1F). We found that Sgo1 N51I mutant that cannot bind Rts1 maintains its ability to interact with condensin (Fig. 1F). Taken together, we have identified a region of Sgo1 _137_LKRTSSRSRSCSLSSPTYSKSYTRLSN_163_ that is required for condensin binding. The identified serine-rich motif (SRM) is essential, but not sufficient for the interaction. Only the fragments that contain in addition the first 100 amino acids of Sgo1, with an extensive coiled coil domain that is required for Sgo1 dimerization (11), can interact with condensin in a pull-down assay. This is consistent with the notion that homodimerization of Sgo1 is a prerequisite of binding to condensin. In contrast, mutant Sgo1 that cannot bind PP2A-Rts1 remains positive for condensin binding *in vitro*. Taken together, the serine-rich motif (SRM) in the N-terminal region of Sgo1 presents a novel functional domain of Sgo1 protein.

### SRM is required for correct chromosome segregation and condensin localization to the centromere

To determine whether the SRM region functions in chromosome segregation, we created a series of deletions of different sizes around the SRM and used the constructs to replace the endogenous *SGO1* allele in haploid yeasts (Fig. 2A). All mutants that lacked the SRM became sensitive to the microtubule depolymerizing drug benomyl to a similar degree as a strain lacking Sgo1. Additionally, all strains lacking the SRM were highly sensitive to the Cik1-cc overexpression that triggers formation of syntelic microtubule–kinetochore attachments at a high frequency (17). In contrast, a deletion of amino acids 201 – 310 did not affect the growth of yeast in the presence of benomyl or upon overexpression of Cik1-cc (Fig. 2B). Additionally, loss of SRM increased chromosome missegregation as monitored by segregation of chromosome IV carrying a *tetO* array and labeled with TetR-GFP (Fig. S2A). The mutants lacking SRM were also not able to maintain the SAC activation and cell cycle arrest in the absence of tension on KTs that was generated by induced loss of sister chromatid cohesion (Fig. S2B). This together demonstrates that the SRM region is required for correct chromosome segregation and centromeric functions as well as for a functional SAC in response to the lack of tension.

**Figure 2.**
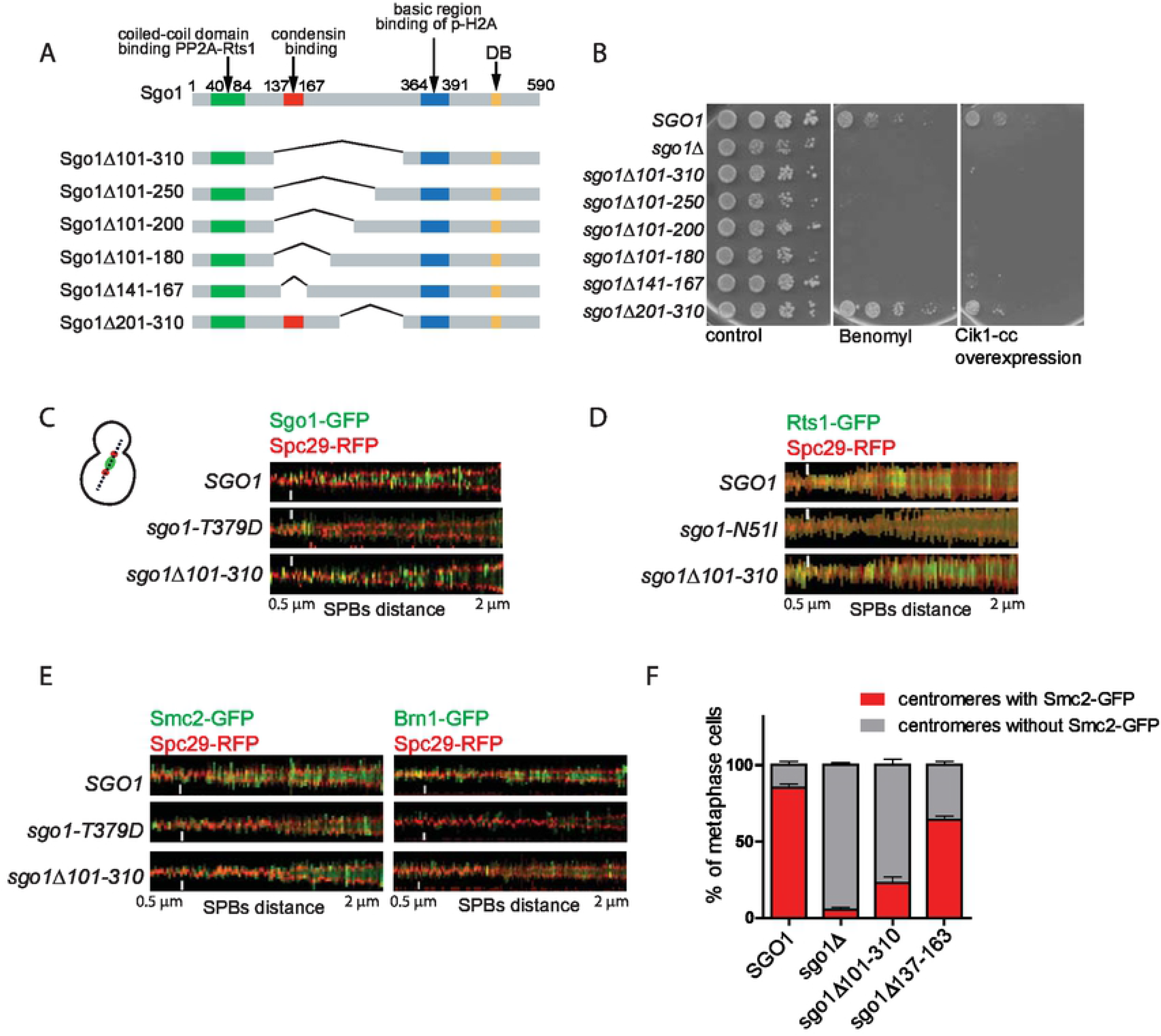
The serine-rich motif (S-rich motif) is required for Sgo1 function in chromosome segregation. A) Schematic depiction of the constructed mutant alleles of *SGO1*. B) Sensitivity of the wild type Sgo1, sgo1Δ and the mutant alleles to the microtubule depolymerizing benomyl (10 μg/ml) and to the overexpression of Cik1-cc that increases the formation of syntelic attachments. C) Signal scans across a line connecting spindle pole bodies (SPBs marked with Spc29-RFP) of single cells were assembled from 150 images to generate a V-plot showing the Sgo1-GFP and Sgo1Δ101-310-GFP localization as the SPBs separation increases. Sgo1-T379D-GFP that fails to localize to the centromeres was used as a control. D) V-plot showing Rts1-GFP localization in wild type cells, or cells carrying Sgo1-N51I (control mutant that fails to localize Rrts1 to the centromere region) and Sgo1Δ101-310 mutations as SPBs distance increases. E) V-plot showing the localization of condensin subunits Smc2-GFP and Brn1-GFP in wild type cells, or cells carrying Sgo1-T379D and Sgo1Δ101-310 mutations as SPBs separation increases. F) Quantification of Smc2-GFP localization. Means with SD of three independent experiments are shown. At least 150 cells were scored in each experiment, only spindles with SPBs separation 0.5 to 2 μm and localizing in the mother cell were evaluated. Scale bar in C, D and E: 1 μm.

If SRM mediates the interaction of Sgo1 with the condensin complex, then its deletion should reduce the accumulation of condensin on centromeres. First, we used spinning disc confocal microscopy to determine whether the mutant Sgo1-GFP lacking SRM localizes to the centromeres. Indeed, the Sgo1 Δ101-301 lacking the SRM localizes between the two spindle poles similarly as wt Sgo1-GFP. This is in a striking contrast to Sgo1 T379D mutant that carries a mutation in the basic region that is responsible for the interaction with the phosphorylated H2A (Fig. 2C). The correct localization of the Sgo1 lacking SRM is further confirmed by the fact that Rts1 localizes to the centromeres in strains carrying the *sgo1 Δ101-310* mutation as efficiently as in the wild type strains (Fig. 2D). Sgo1 also mediates correct localization of Ipl1, the catalytic subunit of the chromosome passenger complex (CPC), to the centromeres. Deletion of SRM or *Δ101-301* did not affect the Ipl1 localization significantly (Fig. S2C,D). In contrast, the fraction of mitotic cells with condensin subunits Smc2-GFP and Brn1-GFP localized to the centromeric region was reduced in yeast strains carrying the *sgo1 Δ101-301* or *sgo1 Δ137-163* mutations (Fig. 2 E, F). The localization of condensin to the rDNA that is observed in budding yeast, was not affected by the mutation in SRM (see below), confirming that the Sgo1-condensin interactions serves only for the enrichment of condensin to centromeres.

### Localization of PP2A and condensin to the centromere are key functions of shugoshin

The evidence so far suggests that Sgo1 serves as a platform for recruitment of Rts1-PP2A and condensin to the centromere. We asked whether these functions are the essential functions of Sgo1 in mitosis. To this end, we created the “Sgo1-mini” protein that consists of the coiled-coil region required for its dimerization and interaction with Rts1 (aa 44 - 84), the SRM important for the recruitment of condensin (aa 137 – 163) and the H2A-binding motif within the basic region that ensures localization of Sgo1 to the centromere (aa 364 – 391). The functional motifs were joined with linkers of 10 amino acids of random sequence. Additionally, we added the C-terminal SV40 nuclear localization signal (NLS) that was previously used to guide Sgo1 to the nucleus (Fig. 3A)(18). The construct “*sgo1-mini*” was further fused with eGFP and integrated at the endogenous locus under the control of the Sgo1 promoter. Live cell imaging revealed that Sgo1-mini localizes to the vicinity of the metaphase spindles, although the localization is somewhat impaired in comparison to the wild type (Fig. 3B). To test whether Sgo1-mini maintains the ability to interact with the PP2A/Rts1 and condensin, a GST-tagged version of Sgo1-mini was over-expressed and purified from *Escherichia coli* BL-21 cells (Fig. S2E) for a GST-pulldown assay and incubated with yeast extracts. This showed that Sgo1-mini interacted with Rts1 and with the condensin subunits Ycg1 and Smc2, although weaker than the full length Sgo1 (Fig. 3C). Remarkably, Sgo1-mini partially rescued the growth defects of *sgo1*Δ cells on benomyl plates as well as the sensitivity of *sgo1*Δ cells to high levels of syntelic attachments caused by Cik1-cc overexpression (Fig. 3D). An equivalent of the N51I mutation that abolishes the interaction with Rts1, and the T379D mutation that impairs centromeric localization of Sgo1, completely abolished the ability of Sgo1-mini to rescue yeast growth under conditions inducing frequent chromosome missegregation (Fig. 3D). Thus, recruitment of the condensin and PP2A to the pericentric region of yeast chromosomes are among the key functions of Sgo1 in mitosis.

**Figure 3.**
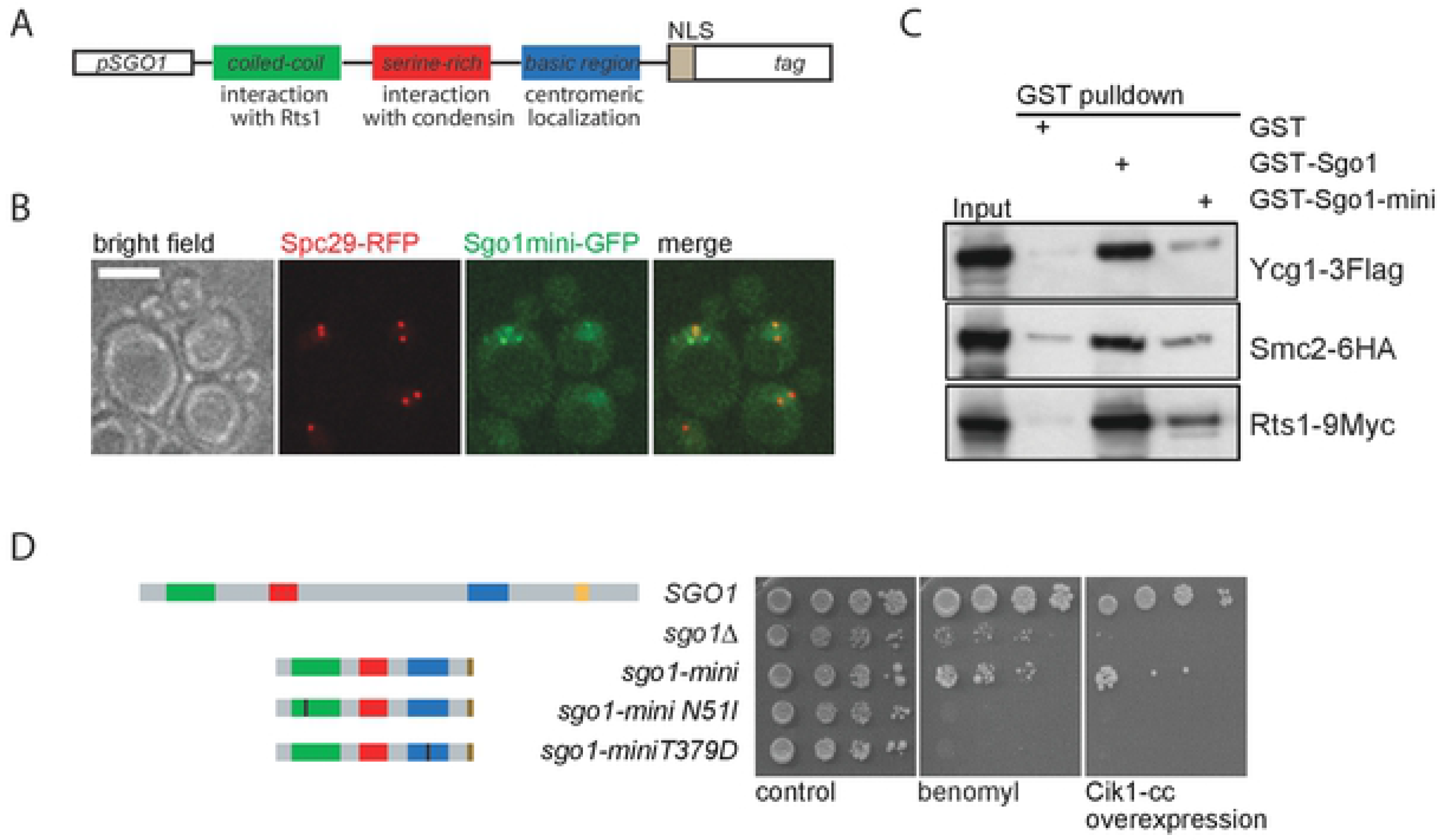
The identified domains support most mitotic functions of Sgo1. A) Molecular organization of Sgo1-mini. B) Sgo1-mini localizes to the nucleus in a close vicinity to the spindle pole bodies (Spc29-RFP). C) Sgo1-mini interacts with condensin and with Rts1 in GST-pulldown experiments. For Coomassie stained gel of purified GST-tagged proteins see Fig. S2E. D) Sensitivity of *sgo1*Δ to the microtubule depolymerizing drug benomyl as well as to Cik1-cc overexpression upon expression of Sgo1-mini, sgo1-mini N51I or sgo1-miniT379D.

### Phosphoregulation of the Sgo1 and condensin interaction

The identified SRM that is required for Sgo1 interaction with condensin is rich in residues that can be modified by phosphorylation. Since yeast Sgo1 is readily phosphorylated (Fig. S3A, B), we asked whether phosphorylation within SRM affects the Sgo1-condensin interaction. Indeed, treatment of cell lysates with phosphatase prior to *in vivo* condensin pulldown increased the amount of recovered Sgo1, suggesting a stronger Sgo-condensin interaction. Strikingly, condensin pulldown from cells harboring *sgo1-N51I* shows diminished interaction (Fig. 4A and (14)), supporting the idea that PP2A-Rts1 activity is crucial for the interaction between Sgo1 and condensin. This is in agreement with our previous finding that centromeric localization of condensin was dramatically lost in these cells (14). To further confirm that the Sgo1 pool that engages condensin at centromeres is in non-phosphorylated form, we isolated the condensin-bound and the condensin-unbound fractions of Sgo1. The fractions were resolved by Phostag^™^ SDS-PAGE to explore their phosphorylation pattern. While the majority of the Sgo1 input and the condensin-unbound Sgo1 appeared to be extensively phosphorylated based on a slow migration in the gel, the condensin-bound Sgo1 exhibited similar migration pattern as a phosphatase treated Sgo1 (Fig. 4B).

**Figure 4.**
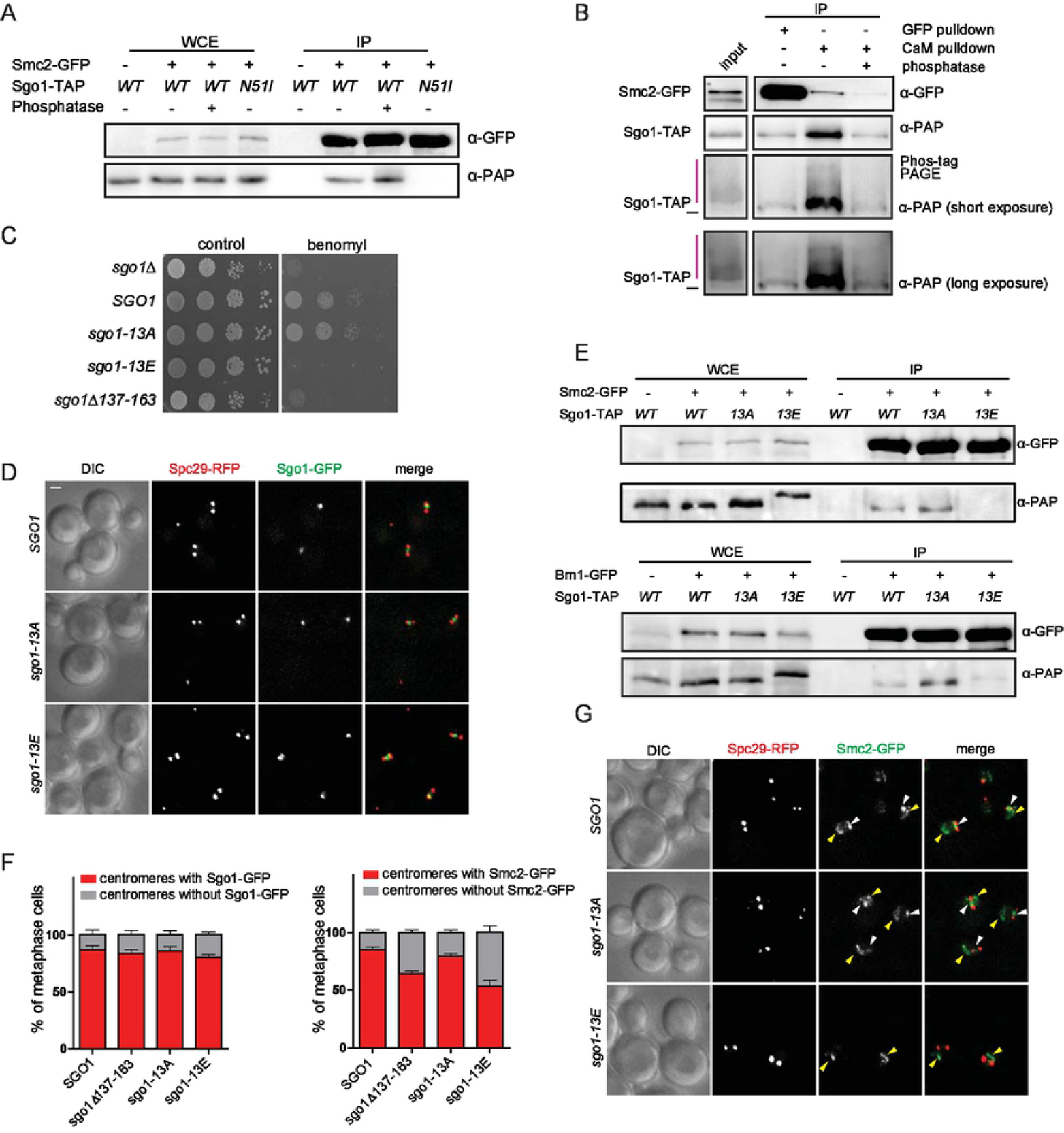
Phosphorylation of the SRM inhibits the binding of Sgo1 to condensin. A) Immunoprecipitation of Smc2-GFP pulled down via the GFP-trap from cell lysates of Sgo1 wt, Sgo1 wt incubated with alkaline phosphatase for 30 minutes at 37°C or Sgo1-N51I. Immunoblots were developed with anti-GFP (Smc2-GFP) and anti-PAP (Sgo1-TAP) antibodies. B) Phosphorylation analysis of Sgo1 bound or unbound to Smc2. Cell lysates from Smc2-GFP, Sgo1-TAP strain were incubated with GFP trap beads to isolate Smc2-GFP and the associated Sgo1. The washout was then incubated with CaM beads to capture the Smc2-unbound Sgo1 fraction; 10% of this fraction was treated with alkaline phosphatase. The red bar marks the phosphoproteins that are migrating slower in the Phostag-PAGE. C) Sensitivity to the microtubule depolymerizing drug benomyl upon expression of SRM-phosphomimicking and phosphoresistant alleles of *sgo1*. D) Localization of *sgo1-13A* and *sgo1-13E*. Spc29-RFP marks the SPBs. E) Immunoprecipitation of Sgo1-TAP with Smc2-GFP or Brn1-GFP pulled down via the GFP-trap. Sgo1wt, Sgo1-13A and Sgo1-13E were analyzed by immunoblotting with anti-GFP (Smc2-GFP, Brn1-GFP, respectively) and anti-PAP (Sgo1-TAP) antibodies. Line 1-4: Input -yeast whole cell extract. F) Evaluation of the Sgo1 and Smc2 centromeric localization in strains carrying the phosphomimicking and phosphoresistant mutant alleles of *SGO1*. G) Localization of Smc2-GFP in cells carrying the phosphomimetic 13E or phosphoreistant 13A mutant alleles. Yellow arrowhead: condensin localized to the rDNA, white arrowhead: condensin localized to pre-anaphase spindles.

Next, we asked whether phosphomimetic or phosphoresistant mutations affect the function of Sgo1. While mutation of all S and T sites to phosphoresistant alanine (*sgo1-13A*) did not affect the sensitivity of yeast cells to benomyl, the strain carrying phosphomimetic mutations (*sgo1-13E*) was highly sensitive to benomyl (Fig. 4C). This was not because of defective Sgo1 localization, as both mutants localized similarly as the wild type protein (Fig. 4D, F) and were expressed to the same level (Fig. S4A). Strikingly, the sgo1-13E failed to interact with condensin in a pulldown experiment (Fig. 4E) and the localization of condensin to the centromere was significantly impaired, while the phosphoresistant mutant showed no discernible phenotype (Fig. 4E-G). Based on these findings, we conclude that the SRM motif of Sgo1 bound to condensin is largely unphosphorylated and that PP2A-Rts1directly or indirectly modulates the phosphorylation state.

To determine whether there are individual sites within the SRM responsible for the phosphoregulation, we performed mass spectrometry of Sgo1 purified from yeast cells arrested in mitosis by the spindle poison nocodazole. This analysis revealed multiple potential phosphorylation sites, among them three sites located within the SRM region, namely S148, S151 and T159, an S421/423 site, and three sites, S487, S518 and S522 near the destruction box (Fig. 5A). To evaluate the function of these sites in chromosome segregation, we constructed a series of yeast strains containing single mutations or their combinations either to phosphoresistant variants with the S/T residues changed to A, or to phosphomimicking variants with the S/T residues changed to E. Phosphomimicking and phosphoresistant mutants of Sgo1 on S421 alone or combined with S423 showed only subtle changes in sensitivity to benomyl or Cik1-cc overexpression and were not further evaluated (data not shown). Mutation of all three sites near the destruction box affected the stability of the Sgo1 protein without altering its localization or localization of condensin nor changed the cellular sensitivity to benomyl (Fig. S4 B-F). This is in line with the previous findings that chromosome segregation in budding yeast mitosis is not substantially affected by the stability and abundance of Sgo1 (13).

**Figure 5.**
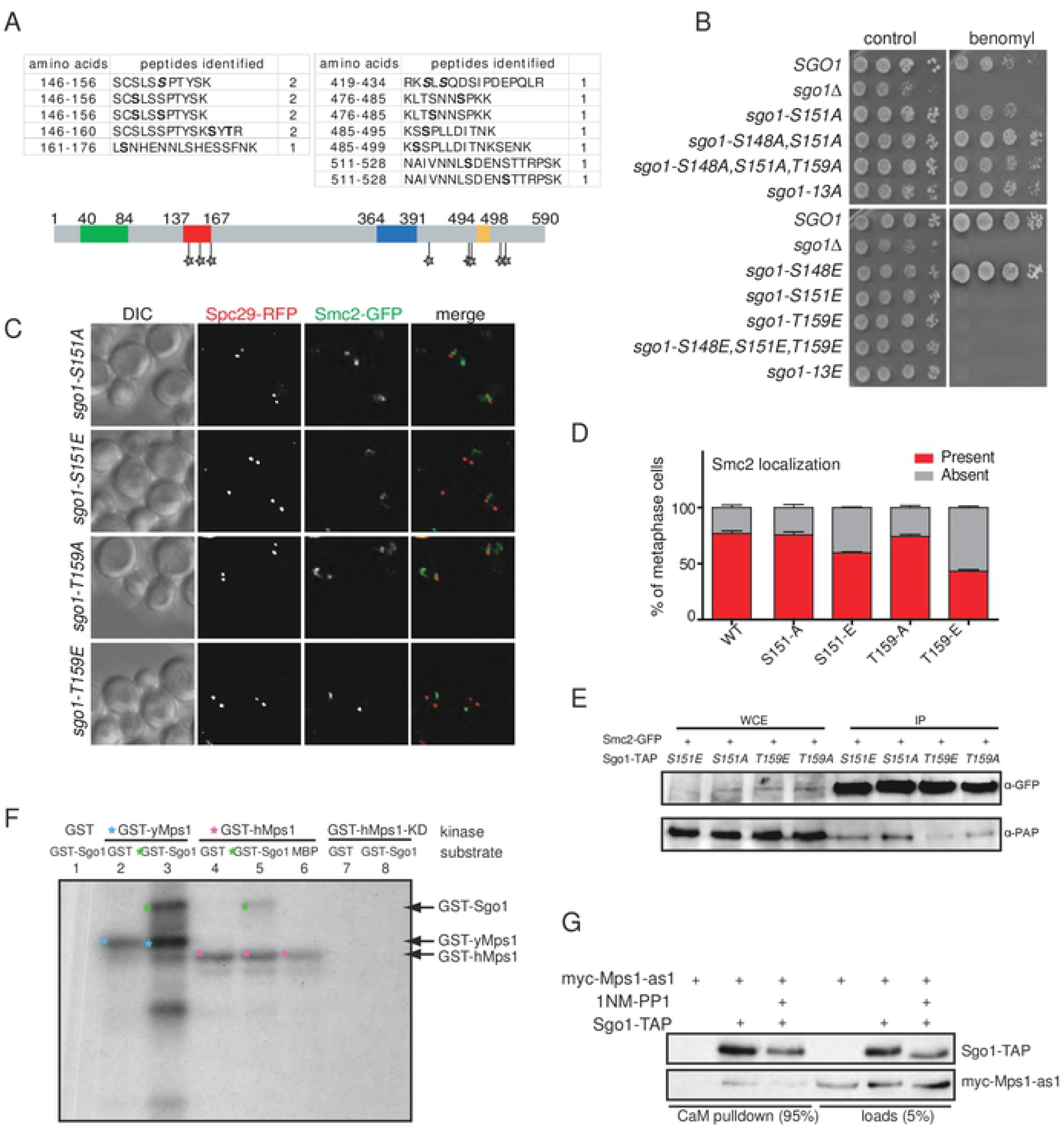
Sgo1-condensin interaction is regulated by phosphorylation. A) Cartoon of Sgo1 highlighting the putative phosphorylation sites on Sgo1 identified by mass spectrometry. B) Sensitivity of the phoshomimicking and phosphoresistant single point mutants alone and in combination to the microtubule depolymerizing drug benomyl. C) Examples of localization of condensin subunits in phosphomimicking and phoshporesistant mutants of Sgo1. D) Quantification of Smc1 localization in the mutants carrying phoshomimicking and phoshoresistant *sgo1* alleles. E) Immunoprecipitation of Sgo1-TAP with Smc2-GFP pulled down via GFP-trap. Sgo1 wt, and the respective phosphomimicking and phosphoresistant mutants were analyzed by immunoblotting with anti-GFP (Smc2-GFP, Brn1-GFP, respectively) and anti-PAP (Sgo1-TAP) antibodies. Line 1-4: Input - yeast whole cell extract. F) *In vitro* kinase assay with purified GST-Mps1 and GST-Sgo1. G) CaM pull down of Sgo1-TAP and myc-Mps1 in budding yeast.

To determine whether there are phosphosites that affect the Sgo1-condensin interaction, we focused on the sites identified within the SRM. Analysis of the phosphoresistant mutants of the sites S148A, S151A and T159A or their combination showed no discernible phenotypes on plates containing benomyl (Fig. 5B). In contrast, the phosphomimicking S151E and T159E markedly increased the sensitivity of yeasts to benomyl, while mutation of S148 did not affect the phenotype (Fig. 5B). The increased benomyl sensitivity is not due to altered Sgo1 structure, because a change of the S/T residues to another bulky or charged amino acid residue did not affect the phenotype (Fig. S4G). Next, we asked whether the mutations of the phosphosites affects the localization of condensin subunits to the centromeric regions. To this end, we analyzed Smc2-GFP localization in Sgo1 phosphomimetic and phosphoresistant mutants. Indeed, Smc2-GFP was mislocalized in a significant fraction of the cells carrying the *sgo1 T159E* mutation and, to a lesser extent, in *sgo1 S151E* mutant (Fig. 5C, D). In contrast, the SMC2-GFP localization was not discernably altered in the phosphoresistant mutants compared to the wild type Sgo1 (Fig. 5C, D). Finally, we tested the interaction between Sgo1 mutants and condensin subunits in a pull down experiment. This experiment revealed that the T159E mutation decreased the ability of Sgo1 to interact with the Smc2-GFP subunit of condensin (Fig. 5E). The interaction between Sgo1 T151E and Smc2-GFP was also partially diminished, although to a lesser extent, reflecting the changes observed in condensin localization (compare Fig. 5C, D, E). Importantly, none of these mutations affected the localization of Sgo1 nor the ability of Sgo1 to interact with Rts1 (Fig. S4H and data not shown). Based on these results we conclude that the interaction of Sgo1 with condensin is regulated by phosphorylation of the SRM region on the S151 and T159 residues.

### Mps1 regulates the Sgo1-condensin interaction through SRM phosphorylation

An open question remains what kinase phosphorylates Sgo1 in the SRM region. We monitored the phosphorylation status of Sgo1 *in vivo* in yeast strains carrying deletions, temperature sensitive or analog sensitive alleles of mitotic kinases Bub1, Ipl1 and Mps1 that are known to affect the Sgo1 function (10, 19-21). While we did not observe any striking changes in the phosphorylation pattern of Sgo1 in the absence of Bub1 and Ipl1 kinase activity, we noticed a loss of phosphorylation upon Mps1 inhibition (Fig. S3 C-G). Therefore, we asked whether Mps1 kinase can phosphorylate Sgo1. By *in vitro* kinase assay we determined that Mps1-GST purified from *E. coli* phosphorylates purified Sgo1-GST. Interestingly, human Mps1-GST also phosphorylates Sgo1 from yeast, while the kinase dead mutant loses this ability (Fig. 5F). Additionally, by immunoprecipitation experiments we determined that in budding yeast Sgo1 interacts with the Mps1 kinase. This interaction is independent of the kinase activity, as inhibition of the kinase activity of Mps1-as (22) by adding the ATP analog 1NM-PP1 did not abolish the interaction (Fig. 5G). Taken together, our data suggest that Sgo1 is directly phosphorylated by Mps1 and this phosphorylation negatively affects the Sgo1-condensin interaction.

## Discussion

### Novel motif within Sgo1 is required for the Sgo1-condensin interaction

Shugoshin proteins have an important role in establishing biorientation during mitosis and meiosis that is executed via coordinated interaction with several proteins at the centromere and the pericentric regions of chromosomes. In budding yeast, Sgo1 also facilitates centromeric localization of condensin and this localization is essential for correct biorientation of sister chromatids (14, 16). This is likely due to the function of condensin, together with cohesin, in establishing the spatial configuration of the pericentric chromatin, which is essential for building functional kinetochore (23). By identification of the interacting region of Sgo1, we were able to create separation-of-function mutations that allowed us to study the consequences of the lack of Sgo1-condensin interaction separately from other Sgo1 functions. Our findings demonstrate that the pericentric enrichment of condensin that is mediated by Sgo1 is required not only for the spatial bias of sister kinetochores that promotes biorientation (16), but also for tension sensing and error correction. Condensin together with cohesin was previously proposed to build a centromeric spring that balances the forces on the metaphase spindle and might be required for tension sensing. Our observations provide additional support for this model (23). Chromosome condensation is also essential for spatial organization and successful partitioning of metaphase chromosomes. Our results further emphasize the importance of Sgo1 for this processes highlighted by recent findings that upon deletion of a centromere from yeast chromosomes, Sgo1 cannot facilitate the condensation of chromosomes (24).

Our data show that the newly identified Serine-Rich Motif of Sgo1 is required for the interaction with condensin and for its localization to the centromeres, while condensin localization to rDNA remains unaffected (Fig. 2E, 4G, S4H). It should be noted that the localization of the condensin at the centromere, while severally impaired, is not completely abolished in the SRM deletion and the 13E mutant. This suggests that there might be another binding site within Sgo1 that was missed in our analysis. For example, the peptide array would not identify interacting domains that are larger than 15 amino acids, or that rely on the tertiary structure of Sgo1. Mutations of Sgo1 that reduce the centromeric localization of condensin lead to defects in chromosome segregation as well as in error correction that leads to SAC activation, and this defect is comparable to a defect observed upon loss of *sgo1* (Fig. 2B, S2A,B). Although we cannot exclude that the loss of SRM does not affect interaction of Sgo1 with other proteins or impairing their activity, our current results suggest that the main defect in SRM is the condensin mislocalization. Therefore, we conclude that the defect in SAC activation is not due to mislocalized CPC with its catalytically active Ipl1 subunit, because lack of SRM does not appear to affect Ipl1 localization. We propose that the correct condensin localization, mediated by its interaction with Sgo1, is required for tension sensing on the MT-KT attachment, most likely via maintaining the spring-like structure of pericentric chromatin that was proposed to contribute to the spindle function (23, 25).

### Mps1 phosporylates Sgo1 to regulate the Sgo1-condensin interaction

Phosphoregulation of the Sgo1-condensin interaction could explain how Sgo1 can facilitates both recruitment of condensin to the centromere in metaphase (14, 16) as well as the spreading of the condensation signal to the chromosome arms in late metaphase (24). Here we demonstrate that Sgo1 interacts with condensin only when the SRM motif remains unphosphorylated (Fig. 4). This unphosphorylated state can be achieved by active dephosphorylation, for example by the interacting phosphatase PP2A (11). In this scenario, PP2A dephosphorylates the SRM within Sgo1, thereby regulating the Sgo1-condensin interaction. Indeed, previous data showed that PP2A-Rts1 is crucial for Sgo1-condensin interaction and as well as condensin localization (14); Verzijlbergen et al., 2014 and this work). However, shugoshins have not been identified as putative substrates in the previous studies of PP2A targets (26, 27). Another possibility is that binding of Sgo1 to condensin or to PP2A protects the SRM from phosphorylation.

An important question arising from our results was the identity of the kinase that phosphorylates the SRM of Sgo1. Using an *in vitro* kinase assay and *in vivo* pull down, we show that the conserved SAC kinase Mps1 interacts with Sgo1 and phosphorylates this protein. Moreover, mass spectrometry determined phosphosites within the SRM motif that were sensitive to the Mps1 activity. This is in agreement with previous studies that suggested that Mps1 kinase directly or indirectly affects the function of Sgo1 (10, 21). We propose that the phosphorylation of Sgo1 by Mps1 might be essential for the release of the condensin load to free the Sgo1 platform to acquire new cargo. In this model, both PP2A-Rts1 and Mps1 fine-tune Sgo1 function and modulate recruitment of proteins required for biorentation. While PP2A phosphatase activity safeguards the efficient Sgo1-condensin interaction, Mps1 kinase activity is required to release the condensin load. We further suggest that the interaction between Sgo1 and condensin on yeast centromeres is the default state during mitosis that becomes disrupted by phosphorylation. Our hypothetical model is that the SRM cannot be phosphorylated when bound to condensin for steric reasons or due to the PP2A-Rts1 mediated dephosphorylation. This model envisions a biorientation machinery where PP2A-Rts1 protection counteracts Mps1 mediated phosphorylation of Sgo1 SRM to orchestrate dynamic condensin loading on the centromere that is essential to create biorientation microenvironment. Increasing tension on the kinetochore and pericentric chromatin may then lead to chromatin stretching, which allows phosphorylation of the SRM. The phosphorylation would then disable the Sgo1-condensin interaction upon successful biorientation and might coincide with the metaphase-to-anaphase transition and Sgo1 removal from centromeres.

## Material and Methods

### Used antibodies

Anti-V5, 1:2000, AbD Serotec (MCA1360GA); α-HA, 1:1000, Santa Cruz Biotechnology (F-7: sc-7392); α-FLAG, 1:1000, SIGMA (monoclonal anti-FLAG M2); α-myc, 1:1000, Santa Cruz Biotechnology (9E10: sc-40); α-GFP, 1:1000, Santa Cruz Biotechnology (B-2: sc-9996); α-Pgk1, 1:10000, Santa Cruz Biotechnology (F-7: sc-7392); α-PAP, HRP-conjugated, 1:2000 Sigma (1291); Secondary α-mouse Polyclonal Goat IgG, 1:5000, R&D SYSTEMS (FAH007).

### Yeast strains and growth

All yeast strains were derived from the genetic background of W303 or BY4743 and are listed in Supplementary Table S1.

### Plasmid construction

All plasmids used in this study are listed in Supplementary Table S2. Sgo1-mini, Sgo1-13A and Sgo1-13E DNA sequences were synthesized at Integrated DNA Technologies (Belgium).

### Yeast cell extract by trichloroacetic acid (TCA)

Exponentially growing yeast cells were harvested, resuspended in 100 μl lysis buffer and incubated 10 min on ice. 40 μl 100% TCA was added and incubated for 10 min on ice. Precipitated proteins were spun down for 10 min at 4°C, 13 000. The pellet was washed with 1 ml ice-cold acetone, dried at 50°C for 5-20 min and resuspended in 50 μl 2x Laemmli buffer. Protein lysates were boiled for 5 min at 96°C.

### Yeast protein extraction for a GST-pulldown

Exponentially growing yeast cells were harvested, washed with cold PBS, and 300 μl yeast lysis buffer was added and resuspended completely. The tube was filled with 0.5 mm ZIRCONIA/SILICA beads (Biospec) and processed in a bead-beater 6x for 1 min and 30 Hz, with a 5 min break in between each beating. The tubes were pierced, placed in a falcon tube and spun down for 2 min at 1000 rpm. The lysate was transferred to a fresh tube, 30 μl NP-40 (10% stock) was added, rotated for 30 min at 4°C and centrifuged for 5 min at 4°C and 5000 rpm. The supernatant was transferred to a fresh tube.

### GST-protein purification from *Escherichia coli* and pull down

IPTG-induced *E. coli* cells were resuspended in 40 ml GST lysis buffer, lysed with high pressure (40 mBar) using the Emulsiflex-C3 (Avestin). 50 μl lysate was mixed with 50 μl 4x Laemmli buffer as a control sample. 2 mM MgCl2, 40 μl Pefabloc (100x) and 4 μl Benzonase were added to the lysate and incubated on ice for 15 min. The lysate was then centrifuged 20 min at 15 000 rpm, 4°C. 1ml pre-washed GST beads (Glutathione-Sepharose 4 Fast Flow HE) were added to the cleaned supernatant and incubated for 2 hours at 4°C on a rotator wheel. The mixture was washed 20x with 20 ml GST washing buffer on a disposable column (QIAGEN) The GST-fused protein was eluted with 1 ml GST elution buffer and the eluted sample was dialyzed overnight at 4°C in 2 l 1x PBS in a Slide-A-Lyzer Dialysis Cassette G2 10 000 MWCO (Thermo Scientific).

100 μl PierceTM Glutathione Magnetic Beads (Thermo Scientific) and 100 μg protein in 500 μl 1x PBS were mixed and rotated for 1 h at 4°C, followed with 3x washing with 600 μl GST washing/binding buffer 10 min and 2x with yeast lysis buffer (5 min) with yeast lysis buffer. Yeast lysate was added to the beads and rotated for 2 h at 4°C. The protein lysate was removed and the beads were washed 2x for 5 min with Washing buffer 1 and 1x for 5 min with Washing buffer 2. The proteins were eluted with 50 μl elution buffer and mixed with 50 μl 4x Laemmli buffer. Samples and the beads were boiled for 5 min at 96°C.

### *In vivo* interaction assay

These experiments were performed as previously described (14).

### GFP Pulldown

Approximately 100 OD_600_ of log-phase cells grown exponentially in YPD were harvested and washed twice with water then lysed by beating with 0.5-mm-diameter glass beads (Sigma-Aldrich) and ice cold, freshly prepared native lysis buffer (20 mM HEPES, pH 7.5, 0.5% Triton X-100, 200 mM NaCl, 1X protease inhibitor mix, 1 mM 1,10 phenanthroline, 1 mM EDTA, 10 mM iodoacetamide). To pulldown GFP-tagged proteins, lysates were incubated for 2 hours at 4°C with 25 μL GFP-Trap coupled agarose beads (Chromotek). After incubation, the agarose beads were washed three times in lysis buffer before samples were eluted with 2X SDS sample buffer. Nitrocellulose membranes were probed with rat anti-GFP (3H9, Chromotek, 1:2,500), PAP (Sigma-Aldrich, 1: 2,500), and mouse anti-HA (Y-11, Santa Cruz Biotechnology, 1:2,500) primary antibodies.

### Phos-tag^™^ SDS-PAGE

Phospho-affinity gel electrophoresis was performed using Phos-tag^™^ acrylamide 7.5% (w/v) running gels polymerized with 50 μM Phos-tag^™^ acrylamide (FUJIFILM Wako Pure Chemical Corporation) and 100 μM MnCl_2_. Gel running and transfer conditions were optimized according to the manufacturer’s protocol.

### Fluorescence microscopy

Exponentially growing yeast cells were imaged by a fully automated Zeiss inverted microscope (AxioObserver Z1) equipped with a MS-2000 stage (Applied Scientific Instrumentation, USA), the CSU-X1 spinning disk confocal head (Yokogawa, Herrsching), LaserStack Launch with selectable laser lines (Intelligent Imaging Innovations, USA) and an X-CITE Fluorescent Illumination System. Images were captured using a CoolSnap HQ camera (Roper Scientific, Canada) under the control of the Slidebook software (Intelligent Imaging Innovations, USA). The fluorescence signal was imaged with a 63x oil objective by using a 473 nm or 561 nm laser. A total of 10 z-stacks were collected at 0.5 μm optical sections. A centromeric localization signal was considered “positive” when the signal intensity on line connecting the two SPBs exceeded the intensity of the average nuclear signal.

### Condensin purification

Condensin was purified from yeast strain YSG0168 (Mat**a**, Smc2-TEVs-HaloTag-TwinStrep::HIS3, Smc4-V5-His6::TRP1). An over-night culture in YPAD medium was inoculated from a single colony. Cells were diluted into 2 L of fresh medium to an OD600 of 0.2. Nocodazole (5 μg/mL final) was added at an OD600 of 0.8. After 1.5 hours of incubation, cells (~12 g) were harvested, washed with PBS twice, resuspended in 12 ml of lysis buffer (150 mM Na phosphate pH 8.0 in 1x PBS, 1x protease inhibitor cocktail (PIC, Sigma)), and flash frozen in liquid nitrogen. Frozen cell suspension was crushed in a swing-mill set to 30 hertz for 3 minutes. After thawing the samples on ice, the cleared supernatant was loaded on 2x 400ul StrepTactin Sepharose (IBA). After washing three times with 2.5 mL of wash buffer (50 mM Na phosphate pH 8.0, 140 mM NaCl, 14 mM Mercaptoethanol). Protein was eluted with 3.2 ml PBS containing 2 mM biotin and 1x PIC.

### Kinase assay

Active or kinase-dead variants of recombinant GST-yMps1 or GST-hMps1 were incubated with 2 μCi [γ-32P] ATP and recombinant GST-Sgo1 in 30 μl of kinase buffer (50 mM Tris pH 7.5, 10 mM MgCl2, 1 mM DTT, 100 μM ATP). Reactions were incubated at 30° C for 30 min and terminated by addition of Laemmli SDS sample dilution buffer and boiling the samples. Proteins were separated on the pre-cast 12.5 % SDS-PAGE (Invitrogen), and phosphorylation was visualized by autoradiography.

### Peptide arrays and far western

Sgo1 peptide arrays were synthesized automatically using SPOT technology with the MultiPep instrument from INTAVIS Bioanalytical Instruments AG as previously (28). To identify the binding sites of Sgo1 for condensin complex protein, 15-mer peptides with 4-mer shift of Sgo1 were synthesized on a membrane. The peptide array membrane was blocked with 5% skim milk in TBST for 1 hour at room temperature, then SV5-PK antibody. The membrane was then washed extensively with TBST before the anti-mouse antibody was added. The membrane was incubated with the antibody at room temperature for another 1 hour prior to standard chemiluminescent detection.

### Identification of the phosphosites by mass spectrometry

To identify the phosphosites on Sgo1, the TAP-tagged protein was overexpressed from a GAL1 promoter for 180 min followed by treatment with nocodazole for 120 min to arrest the overexpressing cells in mitosis. The protein was purified in presence of phosphatases by calmodulin pulldown, eluted with EGTA and separated by PAGE followed by Coomassie staining. The band corresponding to the expected size of Sgo1-TAP was cut from the gel and subjected to the analysis. Mass spectrometry identification of phosphosites on the isolated protein was performed by Nagarunja Nagaraj at the Mass Spectrometry Core Facility at the MPI Biochemistry, Martinsried, Germany, as previously described (29).

## Acknowledgements

We are thankful to Mark Winey and Sabine Elowe for providing plasmids and strains. We thank Jochen Rech (MPI Biochemistry, Martinsried, Germany) for his help with the Sgo1 peptide arrays printing and far western, and Stefan Jentsch for his lasting support during this project. We thank Adele Marston for her kind help with the V-plot analysis and for critical reading of the manuscript. We thank Nagarunja Nagaraj for his help with the identification of the phosphosites on Sgo1 that was performed by the Mass Spectrometry Core Facility at the MPI Biochemistry, Martinsried, Germany. GYM was funded by Georg Forster Research Fellowship awarded by the Alexander-von-Humboldt Foundation. The authors declare no competing financial interests.

## Supporting information legends

**Figure S1 Validation of the Sgo1 - condensin interaction**

(**A**) Far western was performed on a peptide print of Sgo1 using tagged condensin purified from budding yeasts (upper blot) or untagged control. The nonspecific binding is marked in black. Positions A1-A15: peptides covering the Sgo1 sequence from aa 37 to 60 (A9-A13 in the primary screen) and its various mutations. Positions A17-C15: peptides covering the Sgo1 sequence from aa 137 to aa 167 and its various mutations (B15 -B20 in the primary screen). D1 to G20: additional individual peptides that showed week positivity in the primary screen. Positions H17 - H20: tags for antibody controls. (**B**) Comparison of the primary and secondary peptide plot of the putative interacting region aa 37 −60. Upper blot - primary screen, lower blot - secondary screen. Several mutant peptides were added to evaluate the specificity of the putative binding. (**C**) GST-tagged Sgo1 purified from *E. coli*, wt and the mutant variants.

**Figure S2 Recruitment of condensin to the pericentric regions is required for sensing and correction of tensionless attachments**

(A) Chromosome segregation (centromeres of chromosome 4 labeled by tetO/TetR-GFP) in cells released from an arrest in medium containing nocodazole (2 h treatment). Cells were collected 75 min after the releasefrom nocodazole, fixed and imaged. (B) Cell cycle arrest in response to loss of sister chromatid cohesion monitored by immunoblotting of Pds1-9myc degradation in cell lysates. Cells carrying Gal10-Mcd1 allele were arrested in G1 by α-factor and released into medium with and without glucose. Samples were collected at indicated time points. Pgk1 – phosphoglycerate kinase (loading control). (C) Examples of Ipl1-GFP localization in *sgo1*Δ, wild type *SGO1, sgo1Δ137-163* and *sgo1Δ101-310* and (D) its quantification. (E) Purified GST-tagged Sgo1wild type and Sgo1-mini.

**Figure S3 Phosphoregulation of Sgo1 by mitotic kinases**

(A) Migration of Sgo1-TAP in Phostag-PAGE. Cells were collected at specified time points after a release from a G1 arrest by α-factor. After 45 min, α-factor was again added to limit the cells to one cell cycle. (B) Migration of Sgo1-TAP in Phostag-PAGE after release from α-factor. The sample 75/λ was treated with λ phosphatase for 60 min. (C) Migration of Sgo1-TAP in Phostag-PAGE showed similar pattern in the presence or absence of BUB1. (D, E) Migration of Sgo1 in Phostag-PAGE was not affected by the presence or absence of IPL1 kinase activity; in contrast, H3 phosphorylation was almost completely abolished in cells expressing *ipl1-321*, a temperature-sensitive mutation that impairs the kinase activity at 37°C. (F) Migration of Sgo1-TAP extracted from cells harboring an analog sensitive allele of MPS1 *(mps1-as1)* in Phostag-PAGE in presence or absence of the ATP analog 1NM-PP1. (G) Autophosphorylation of Mps1 was monitored in the same experiment as an internal control for kinase activity. * represents phosphorylated form of the indicated proteins.

**Figure S4 Phenotype of Sgo1-phosphomutants**

(**A**) Levels phosphomimicking (13E) and phosphoresistant (13A) mutants of Sgo1 - immunoblot and their quantification. (**B**) Localization of Sgo1-GFP-3A and 3E to preanaphase spindle. (**C**) Localization of condensin in Sgo1-3A and Sgo1-3E mutants. (**D**) Sensitivity of *sgo1-3A* and *sgo1-3E* mutants to benomyl. (**E, F**) Protein levels of Sgo1-3A and Sgo1-3E - immunoblot and quantification. (**G**) Sensitivity of T159 mutants to benomyl. (**H**) Localization of the Sgo1-T159 and Sgo1-S151 mutants to the preanaphase spindle.

### Tables

**Supplementary Table 1 Yeast strains used and created in this study**

**Supplementary table 2 List of plasmids created and used in this work**

## References

1. Indjeian VB, Murray AW. Budding yeast mitotic chromosomes have an intrinsic bias to biorient on the spindle. Curr Biol. 2007;17(21):1837–46.

2. Magidson V, Paul R, Yang N, Ault JG, O’Connell CB, Tikhonenko I, et al. Adaptive changes in the kinetochore architecture facilitate proper spindle assembly. Nat Cell Biol. 2015;17(9):1134–44.

3. Godek KM, Kabeche L, Compton DA. Regulation of kinetochore-microtubule attachments through homeostatic control during mitosis. Nat Rev Mol Cell Biol. 2015;16(1):57–64.

4. Musacchio A. The Molecular Biology of Spindle Assembly Checkpoint Signaling Dynamics. Curr Biol. 2015;25(20):R1002–18.

5. Marston AL. Shugoshins: tension-sensitive pericentromeric adaptors safeguarding chromosome segregation. Mol Cell Biol. 2015;35(4):634–48.

6. Kitajima TS, Kawashima SA, Watanabe Y. The conserved kinetochore protein shugoshin protects centromeric cohesion during meiosis. Nature. 2004;427(6974):510–7.

7. McGuinness BE, Hirota T, Kudo NR, Peters JM, Nasmyth K. Shugoshin prevents dissociation of cohesin from centromeres during mitosis in vertebrate cells. PLoS Biol. 2005;3(3):e86.

8. Tang Z, Shu H, Qi W, Mahmood NA, Mumby MC, Yu H. PP2A is required for centromeric localization of Sgo1 and proper chromosome segregation. Dev Cell. 2006;10(5):575–85.

9. Kawashima SA, Yamagishi Y, Honda T, Ishiguro K, Watanabe Y. Phosphorylation of H2A by Bub1 prevents chromosomal instability through localizing shugoshin. Science. 2010;327(5962):172–7.

10. Storchova Z, Becker JS, Talarek N, Kogelsberger S, Pellman D. Bub1, Sgo1, and Mps1 mediate a distinct pathway for chromosome biorientation in budding yeast. Mol Biol Cell. 2011;22(9):1473–85.

11. Xu Z, Cetin B, Anger M, Cho US, Helmhart W, Nasmyth K, et al. Structure and function of the PP2A-shugoshin interaction. Molecular cell. 2009;35(4):426–41.

12. Nerusheva OO, Galander S, Fernius J, Kelly D, Marston AL. Tension-dependent removal of pericentromeric shugoshin is an indicator of sister chromosome biorientation. Genes Dev. 2014;28(12):1291–309.

13. Eshleman HD, Morgan DO. Sgo1 recruits PP2A to chromosomes to ensure sister chromatid bi-orientation during mitosis. J Cell Sci. 2014;127(Pt 22):4974-83.

14. Peplowska K, Wallek AU, Storchova Z. Sgo1 regulates both condensin and Ipl1/Aurora B to promote chromosome biorientation. PLoS Genet. 2014;10(6):e1004411.

15. Campbell CS, Desai A. Tension sensing by Aurora B kinase is independent of survivin-based centromere localization. Nature. 2013;497(7447):118–21.

16. Verzijlbergen KF, Nerusheva OO, Kelly D, Kerr A, Clift D, de Lima Alves F, et al. Shugoshin biases chromosomes for biorientation through condensin recruitment to the pericentromere. Elife. 2014;3:e01374.

17. Jin F, Liu H, Li P, Yu H-G, Wang Y. Loss of Function of the Cik1/Kar3 Motor Complex Results in Chromosomes with Syntelic Attachment That Are Sensed by the Tension Checkpoint. PLOS Genetics. 2012;8(2):e1002492.

18. Vaur S, Cubizolles F, Plane G, Genier S, Rabitsch PK, Gregan J, et al. Control of Shugoshin function during fission-yeast meiosis. Curr Biol. 2005;15(24):2263–70.

19. Tanno Y, Kitajima TS, Honda T, Ando Y, Ishiguro K, Watanabe Y. Phosphorylation of mammalian Sgo2 by Aurora B recruits PP2A and MCAK to centromeres. Genes Dev. 2010;24(19):2169–79.

20. Liu H, Rankin S, Yu H. Phosphorylation-enabled binding of SGO1-PP2A to cohesin protects sororin and centromeric cohesion during mitosis. Nat Cell Biol. 2013;15(1):40–9.

21. Arguello-Miranda O, Zagoriy I, Mengoli V, Rojas J, Jonak K, Oz T, et al. Casein Kinase 1 Coordinates Cohesin Cleavage, Gametogenesis, and Exit from M Phase in Meiosis II. Dev Cell. 2017;40(1):37–52.

22. Jones MH, Huneycutt BJ, Pearson CG, Zhang C, Morgan G, Shokat K, et al. Chemical genetics reveals a role for Mps1 kinase in kinetochore attachment during mitosis. Curr Biol. 2005;15(2):160–5.

23. Stephens AD, Haase J, Vicci L, Taylor RM, 2nd, Bloom K. Cohesin, condensin, and the intramolecular centromere loop together generate the mitotic chromatin spring. J Cell Biol. 2011;193(7):1167–80.

24. Kruitwagen T, Chymkowitch P, Denoth-Lippuner A, Enserink J, Barral Y. Centromeres License the Mitotic Condensation of Yeast Chromosome Arms. Cell. 2018;175(3):780–95 e15.

25. Stephens AD, Haggerty RA, Vasquez PA, Vicci L, Snider CE, Shi F, et al. Pericentric chromatin loops function as a nonlinear spring in mitotic force balance. J Cell Biol. 2013;200(6):757–72.

26. Zapata J, Dephoure N, Macdonough T, Yu Y, Parnell EJ, Mooring M, et al. PP2ARts1 is a master regulator of pathways that control cell size. J Cell Biol. 2014;204(3):359–76.

27. Wang X, Bajaj R, Bollen M, Peti W, Page R. Expanding the PP2A Interactome by Defining a B56-Specific SLiM. Structure. 2016;24(12):2174–81.

28. Brandt O, Feldner J, Stephan A, Schroder M, Schnolzer M, Arlinghaus HF, et al. PNA microarrays for hybridisation of unlabelled DNA samples. Nucleic Acids Res. 2003;31(19):e119.

29. Gnad F, de Godoy LM, Cox J, Neuhauser N, Ren S, Olsen JV, et al. High-accuracy identification and bioinformatic analysis of in vivo protein phosphorylation sites in yeast. Proteomics. 2009;9(20):4642–52.

